# Assessing allele specific expression across multiple tissues from RNA-seq read data

**DOI:** 10.1101/007211

**Authors:** Matti Pirinen, Tuuli Lappalainen, Noah A. Zaitlen, GTEx Consortium, Emmanouil T. Dermitzakis, Peter Donnelly, Mark I. McCarthy, Manuel A. Rivas

## Abstract

**Motivation:** RNA sequencing enables allele specific expression (ASE) studies that complement standard genotype expression studies for common variants and, importantly, also allow measuring the regulatory impact of rare variants. The Genotype-Tissue Expression project (GTEx) is collecting RNA-seq data on multiple tissues of a same set of individuals and novel methods are required for the analysis of these data.

**Results:** We present a statistical method to compare different patterns of ASE across tissues and to classify genetic variants according to their impact on the tissue-wide expression profile. We focus on strong ASE effects that we are expecting to see for protein-truncating variants, but our method can also be adjusted for other types of ASE effects. We illustrate the method with a real data example on a tissue-wide expression profile of a variant causal for lipoid proteinosis, and with a simulation study to assess our method more generally.

**Availability:** MAMBA software: http://birch.well.ox.ac.uk/∼rivas/mamba/ R source code and data examples: http://www.iki.fi/mpirinen/

## 1 Introduction

Advancements in sequencing technologies are enabling unprecedented possibilities to study the transcriptome. In RNA-sequencing studies, it is possible to distinguish between transcripts from the two haplotypes of an individual using heterozygous sites. This approach, called allele specific expression (ASE) analysis, allows an alternative way to quantify cis-regulatory variation, complementary to eQTL analysis. Additionally, ASE has been utilized to analyze transcriptome effects of nonsense-mediated decay triggered by predicted loss-of-function variants (MacArthur *et al.*, 2012; Montgomery *et al.*, 2011; Lappalainen *et al.*, 2013).

The Genotype-Tissue Expression (GTEx) project is establishing a resource database and tissue bank for the scientific community to study the relationship between genetic variation and gene expression in human tissues (GTEx-Consortium, 2013), with an aim to interpret GWAS findings for translational research. The project is analyzing gene expression from various perspectives, including transcript structure, expression quantity and diversity, eQTLs, and allele specific expression differences. Furthermore, as medical genetics pursues exploration of rare variants, insights gained from the study of DNA and RNA sequencing data in the GTEx project will become important for functional interpretation of rare variants (Rivas *et al.*, 2011, 2013; Zuk *et al.*, 2014).

To date, some methods have been proposed for the analysis of allele specific expression data, but these methods largely focus on a single tissue (Ronald *et al.*, 2005; Zhang *et al.*, 2009; Degner *et al.*, 2009; Sun, 2012) although some could be applied also to multiple tissues (Skelly *et al.*, 2011). The multi-tissue aspect is important for interpreting disease association findings (Grundberg *et al.*, 2012; McCarroll *et al.*, 2008) since eQTL and early ASE studies suggest widespread tissue specific effects of cis-regulatory variants (Dimas *et al.*, 2009; Gutierrez-Arcelus *et al.*, 2013). Currently, more sophisticated methods for cross-tissue eQTL analysis are emerging (Flutre *et al.*, 2013). However, eQTL analysis requires large sample sizes, while ASE analyses can be conducted with significantly smaller data sets.

In this paper we present novel statistical methods for analyzing ASE patterns from RNA-seq data across multiple tissues. Our main motivation are phenomena such as nonsense-mediated decay that are expected to lead to strong ASE where one of the alleles is expressed only a little if at all. From a statistical point of view an advantage of strong ASE is that it can be studied even with a single individual and without very large read counts. (See Middle panel of Fig. 4 for an example data set.)

We address three main questions:

- In which tissues does a heterozygous site show ASE?
- Which tissues show similar ASE effects at the site studied?
- What proportion of a certain class of variants (such as, e.g., protein truncating variants) show ASE in all tissues, only in some tissues, or in no tissues?

For the first question, a standard frequentist version of the binomial test is commonly used. However, an interpretation of such a test depends on factors like the read count and simultaneous analysis of multiple tissues is very challenging. Hence, we require that our statistical framework allows a simultaneous comparison between several cross-tissue models for observed data. For example, we want to weigh relative support of the model where all tissues show ASE to the models where only a single tissue shows ASE and to the null model where none of the tissues show ASE. For this purpose, we adopt a Bayesian model comparison framework. Among its favorable properties are natural ways to compare models with differing number of parameters and to fully account for the amount of available data when evaluating relative support of the models.

## 2 Methods

### 2.1 Grouped tissue model (GTM)

We consider RNA-seq read counts overlapping a particular genomic position from multiple tissues of one individual who is heterozygous at that position. (See the Discussion and Supplementary Information for extensions to multiple sites and individuals.) For tissue *s* = 1, …, *T*, let *y_s_*_1_ and *y_s_*_2_ be the number of reads supporting the reference and non-reference allele, respectively, and let *n_s_* = *y_s_*_1_ + *y_s_*_2_. We classify the tissues into three groups: no ASE (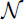), moderate ASE (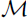) and strong ASE (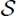) and denote the group of tissue *s* by 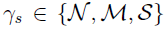. (See Middle panel of Fig. 4 for an example data set that motivates us to use the three groups chosen.) For each group 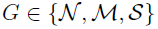, we denote the proportion of the transcripts with the reference allele by *θ*(*G*). We use a binomial sampling model for the data conditional on the group indicators:

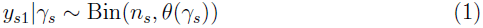

Possible expression states of the tissues differ in the prior assumptions about parameters *θ*(*·*). We use the following priors (Fig. 1) to describe different groups:

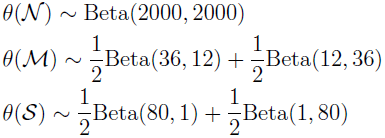

**Figure 1:**
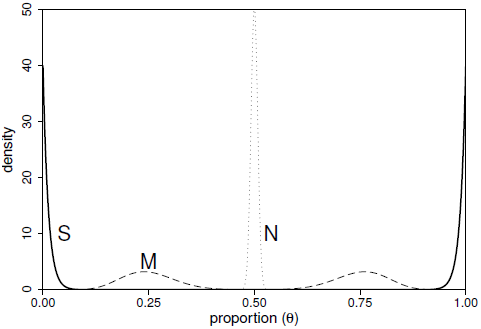
Densities of the prior distributions for the proportion of reference allele for the three groups: 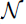, 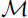, and 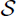.

Under no ASE model 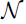 both alleles are expressed (almost) equally and hence *θ*(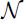) *≈* 0.5. As in Skelly *et al.* (2011), our 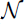 model allows small deviations from exact point value of 0.5 in order to be robust against some technical measurement bias as well as very small ASE effects that are not a main focus in this study. The parameters for Beta distributions have been chosen in a way that clearly separates the three groups from each other (Fig. 1) and thus gives an informative framework to classify the tissues into three groups (see the Supplementary Information for further discussion on the prior specification and extensions to truncated priors and independence across tissues). Our implementation allows user to easily modify the prior parameters as well as use one-sided versions of the prior distributions instead of the two-sided versions used in our simulation experiments. For example, when studying nonsense-mediated decay, we may require that the reference allele is expressed more strongly, and hence consider only the one-sided ASE states. On the other hand, the two-sided ASE states are tailored for the situations where we do not want to make an assumption about which allele might be dominating. This is useful, for example, when studying imprinting.

#### Configurations

For fixed prior distributions, the model space consists of 3*^T^* configurations represented by 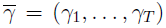 vector where each tissue specific indicator 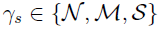.

We partition the space of configurations into five ASE states:

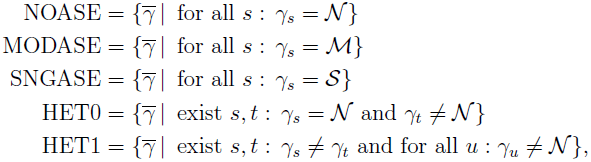

where the states represent configurations for no ASE (NOASE), moderate ASE (MODASE), strong ASE (SNGASE), heterogeneity with at least one tissue showing no ASE (HET0) and heterogeneity with all tissues showing some ASE (HET1). We also consider a tissue specific sub state of the heterogeneity states:

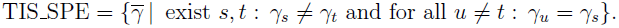

In order to do probabilistic comparison between the states we need to define prior probabilities for each state. We do this by defining prior for each configuration in a way which depends on its distance from homogeneity. We define a distance 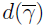 for each configuration 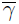 as the smallest number of tissues whose group indicators need to be changed in order to turn 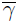 into a homogeneous configuration (either NOASE, MODASE or SNGASE). Formally, 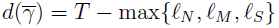, where 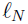, 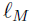 and 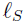 are the number of tissues that 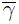 assigns to 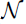, 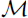, and 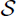, respectively. In particular, the configurations with *d* = 0 are the three homogeneous configurations and the configurations with *d* = 1 form the set TIS_SPE.

We specify the total prior probability for each possible value of *d* = 0, …, *T −* ⌈*T*/3⌉ and then distribute it equally among the configurations with the same distance. This prior allows us to easily implement the idea that among the vast space of configurations we favor *a priori* the parsimonious ones, i.e., those where many tissues are similar. Our prior distribution extends the one recently used for muti-tissue eQTL setting (Flutre *et al.*, 2013) to the case of more than two expression states. In the results reported in this work, we have set a prior mass of 0.75 to *d* = 0 (i.e., 0.25 for each of NOASE, MODASE and SNGASE) and the remaining 0.25 has been divided equally among all possible values of *d* = 1, …, *T −* ⌈*T*/3⌉. This choice was made because it gives an equal prior weight for the four main patterns of ASE: three homogeneous states and general heterogeneity.

For settings where many variants are available for a joint analysis, we extend GTM to a hierarchical model GTM* that learns from the data the proportion of variants belonging to each of the five states and thus avoids the need to fix prior probabilities of the states. We describe GTM* and compare it to GTM in the Supplementary Information.

### 2.2 Computation

We use a standard Gibbs sampler to explore the posterior distribution of the configuration indicators 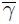 under GTM (see the Supplementary Information for details).

The basic building block for model comparison is the Bayes factor between configuration 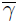 and the NOASE state

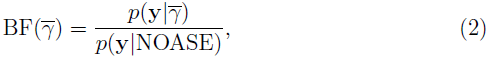

where **y** = (*y*_1*·*_, …, *y_T ·_*). Evaluation of BF(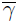) can be done analytically by using Beta-binomial likelihood separately for the numerator and the denominator. Thus we can quickly evaluate the Bayes factor for any particular configuration and compare even hundreds of configurations. However, when the number of tissues is large (say *T >* 10), the number of possible configurations grows too large to be exhaustively evaluated (analogous to a problem with eQTLs in Flutre *et al.* (2013)). This becomes problematic in particular when assessing heterogeneity (either HET0 or HET1), which in principle would require a consideration of all those groupings that assume differences between some tissues. Therefore, we introduce an approximation that avoids enumerating all groupings when assessing heterogeneity, by focusing on only configurations that are strongly supported by the data. Thus we assume that the configurations with *d >* 1 that have not been visited by the Gibbs sampler and are not among a few dozen top heterogeneity models defined by tissue specific group membership probabilities, have negligible marginal likelihood and can be ignored. This leads to a lower bound for the marginal likelihoods of the heterogeneity states. In practice, a comparison to the exact values for cases where exact values can be calculated (*T ≤* 10) shows that the lower bound is actually a good estimate in most cases (see Results).

By combining the Bayes factors with the prior probabilities of the states we have the posterior probabilities of the states.

### 2.3 Q-statistic for heterogeneity

We compare GTM to a standard heterogeneity measure

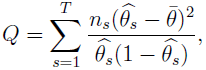

where 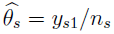 and 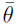 are the empirical proportion of the reference allele at tissue *s* and across all tissues, respectively. (If *y_s_*_1_ = 0, we set *y_s_*_1_ = 0.5 and *y_s_*_2_ = *n_s_ −* 0.5 to avoid numerical problems.) The idea is that this measure increases with the heterogeneity of the tissue specific *θ_s_* parameters, and can thus be used as a measure of heterogeneity. An empirical P-value of the Q-statistic is estimated by simulating data sets where the number of tissues and reads match with the observed data and where all tissues have the same value for *θ_s_* = *θ̄*.

## 3 Results

We first assess performance of GTM on simulated data and compare it with a standard heterogeneity measure. Second, we illustrate GTM on a real data example taken from the Genotype-Tissue Expression (GTEx) project. Results of GTM* are presented in the Supplementary Information.

**Data simulation.** For three values for the number of tissues (*T* = 5, 10, 30) and two values for the number of reads per each tissue (*n_s_* = 10, 50, for all *s*), we simulated 1,000 data sets for nine scenarios given in Table 1. The reference allele read count for each tissue *s* was sampled from Bin(*n_s_, θ_s_*), where *θ_s_* is 0.5 for group 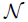, 0.75 for group 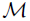 and 0.99 for group 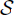.

**Table 1:** Scenarios for simulations.

| | $\mathcal{N}$ | $\mathcal{M}$ | $\mathcal{S}$ | STATE |
| --- | --- | --- | --- | --- |
| 1 | $T$ | 0 | 0 | NOASE |
| 2 | 0 | $T$ | 0 | MODASE |
| 3 | 0 | 0 | $T$ | SNGASE |
| 4 | $T - 1$ | 0 | 1 | HET0 |
| 5 | 1 | 0 | $T - 1$ | HET0 |
| 6 | 0 | $T - 1$ | 1 | HET1 |
| 7 | 0 | 1 | $T - 1$ | HET1 |
| 8 | $\lceil T/2 \rceil$ | $\lfloor T/2 \rfloor$ | 0 | HET0 |
| 9 | 0 | $\lceil T/2 \rceil$ | $\lfloor T/2 \rfloor$ | HET1 |

For each of the nine scenarios, Table shows how many tissues (out of total $T$ ) belong to each of the three possible groups, $\mathcal{N}$ , $\mathcal{M}$ and $\mathcal{S}$ , whose $\theta$ parameters are 0.5, 0.75 and 0.99, respectively. The STATE column shows to which of the five combined states each scenario belongs.

**GTM results.** We applied GTM on the simulated data sets with 2,000 Gibbs sampler iterations and show results in Figure 2. The run time was 8, 16 and 53 seconds per data set, for *T* = 5, 10 and 30, respectively (Intel i7 /3.40 GHz).

**Figure 2:**
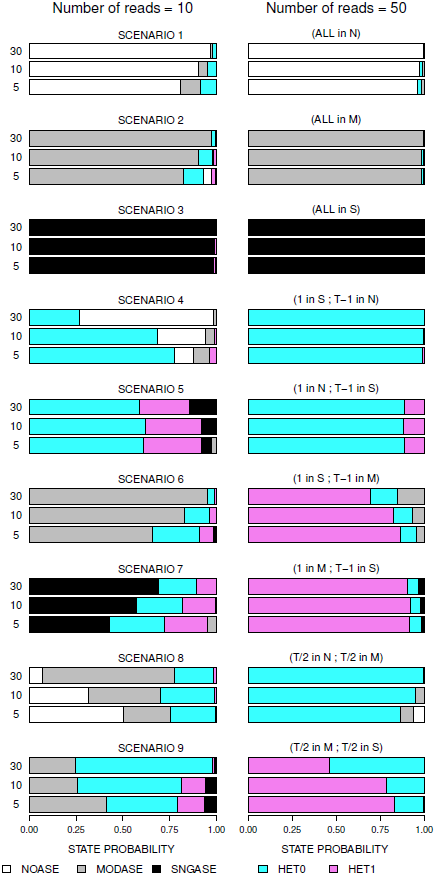
Each of the nine simulation scenarios (Table 1) is represented by three numbers of tissues (5, 10, 30) and two values for number of reads (10, left column and 50, right column). Each bar is divided into five colors (map given at the bottom) according to the (average) posterior probability of the five states when GTM was applied to the simulated data sets.

For the homogeneous scenarios 1, 2 and 3, the detectability of the true state increases with the number of tissues, and in general is already high with only 5 tissues and 10 reads per each tissue. In particular, when the true state is SNGASE (scenario 3), there is no noticeable uncertainty about the correct state in any of the data sets, while for NOASE (scenario 1) and MODASE (scenario 2) the data are not equally decisive. This is related to the fact that the variance in the read counts is larger for *θ* = 0.5 (NOASE) and *θ* = 0.75 (MODASE) than it is for the extreme value of *θ* = 0.99 (SNGASE).

Our scenarios 4, 5, 6 and 7 consider the smallest possible amount of heterogeneity: a single tissue is different from the others. In these scenarios we see an opposite trend from the homogeneous ones: the true heterogeneous state becomes harder to distinguish from the closest homogeneous one as the number of tissues increases. This behavior implements an idea that if one tissue seems to deviate from the others, we believe more in that distinction if only a couple of other tissues are analyzed, than if there are tens of other tissue, that are all similar to each other. Information about the true state increases with the number of reads per tissue, and the overall heterogeneity probability (HET0 + HET1) is high for all heterogeneous scenarios for read count 50.

The scenarios 8 and 9 represent stronger heterogeneity where about half of the tissues belong to one group and the remaining half to another. When the true underlying groups are 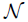 and 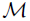 (scenario 8), then 10 reads is not yet enough to clearly separate the true pattern from the homogeneous states (NOASE and MODASE), while 50 reads is enough for this purpose. In the scenario 9, where tissues are divided between 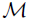 and 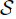, the overall heterogeneity probability is always fairly large, but it is difficult to exclude the possibility that at least one tissue belongs to 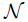, especially when many tissues are available. As a consequence, the HET0 state gets a considerable probability even though the true underlying state is HET1. To distinguish between the two heterogeneity states in this scenario requires read counts larger than 50.

Taken together, the results in Figure 2 show that the model is correctly able to distinguish between all five combined states but an amount of information required for accurate classification varies between scenarios.

**Comparisons with Q-statistic.** To show differences between our GTM heterogeneity probability and Q-statistic in detecting heterogeneity we show ROC curves for two settings (Fig. 3). In the first one, we use the 1,000 data sets from simulation scenario 1 to represent a homogeneous state and the 1,000 data sets from scenario 4 to represent a heterogeneous state, with *T* = 30 tissues and *n* = 10 reads per tissue. The ROC curve on the left panel of Figure 3 shows how those 2,000 data sets are ranked by the posterior probability of HET0+HET1 state from GTM and by the empirical P-value of Q. The right panel in Figure 3 shows similar results when comparing the homogeneous scenario 3 to the heterogeneous scenario 7. In both settings GTM is slightly better in detecting heterogeneity than the Q-statistic: ROC curve of GTM is consistently above that of Q. We discuss the difference between the two approaches in Discussion.

**Figure 3:**
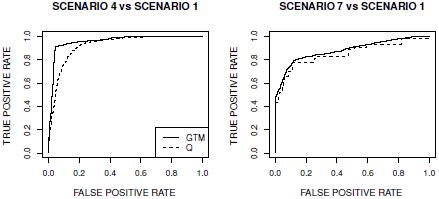
ROC curves for detecting heterogeneity using GTM and Q-statistic on simulated data sets. Left: scenario 1 (homogeneity) vs scenario 4 (heterogeneity). Right: scenario 3 (homogeneity) vs scenario 7 (heterogeneity). Parameters are *T* = 30 tissues and *n* = 10 reads per tissue. Heterogeneity statistics are the posterior probability of HET0+HET1 from GTM and the empirical P-value from the Q-statistic.

**Accuracy of approximation.** To assess how accurate our approximation for marginal likelihoods of the heterogeneity states is, we compared the posterior probabilities of the five states from GTM with the exact values on all 9,000 data sets simulated with *T* = 10 tissues and *n* = 10 reads per tissue. As a measure of accuracy we use the total variation distance (TV), which for discrete distributions (*p_i_*) and (*q_i_*) is defined as 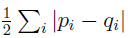. TV describes how much of the probability mass needs to be relocated in order to turn the first distribution into the other, and also gives an upper bound for the maximal difference in probability that the two distributions assign to any one state.

We observed (Table 2) that 95% of the data sets had TV *<* 0.05, and 0.5% had TV *>* 0.10, with maximal TV being 0.168. As TV less than 0.10 is unlikely to change our inference on the underlying state, our approximation works well for a great majority of the data sets tested. However, there are large differences in the accuracy between the scenarios with scenario 8 (5 tissues in 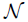 and other 5 in 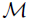) showing the strongest discrepancy. An explanation for this is that with only 10 reads per tissue the scenario 8 assign non-negligible posterior probability to so many of the configurations showing heterogeneity between 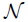 and 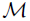 groups that some of them are missed during our default number of 2,000 Gibbs sampler iterations.

**Table 2:** Accuracy of GTM.

| SCENAR | TV < 0.01 | TV < 0.05 | TV < 0.10 | max |
| --- | --- | --- | --- | --- |
| 1 | 0.462 | 0.972 | 1 | 0.092 |
| 2 | 0.452 | 0.959 | 0.995 | 0.129 |
| 3 | 1 | 1 | 1 | 0.006 |
| 4 | 0.160 | 0.986 | 0.999 | 0.105 |
| 5 | 0.997 | 1 | 1 | 0.011 |
| 6 | 0.327 | 0.973 | 1 | 0.070 |
| 7 | 0.994 | 1 | 1 | 0.012 |
| 8 | 0.081 | 0.663 | 0.962 | 0.168 |
| 9 | 0.565 | 1 | 1 | 0.025 |

For each of the nine scenarios, Table shows the proportion of simulated data sets (out of total 1,000 with $T = 10$ tissues and $n = 10$ reads per tissue) for which total variation (TV) distance between GTM estimates and the exact posterior distribution of the five states is less than 0.01, 0.05 or 0.10. The max column shows the maximum TV distance observed.

In Discussion we propose an additional heterogeneity measure to complement our approximation for the posterior probability of heterogeneity states, especially for data sets with large number of tissues.

### 3.1 A protein truncating variant in *ECM1*

We consider read count data on SNP rs121909115 in chromosome 1, whose non-reference allele (T) introduces a stop codon in some transcripts of the gene *ECM1*. This is an example of a protein-truncating mutation that we expect to experience nonsense-mediated decay (NMD) leading to a reduction in the transcripts from the non-reference allele. However, the strength of NMD and its consistency across different tissue types is unknown. In the currently available GTEx data we have one individual who is heterozygous at this SNP and Figure 4 shows the data and GTM results across 7 tissues.

**Figure 4:**
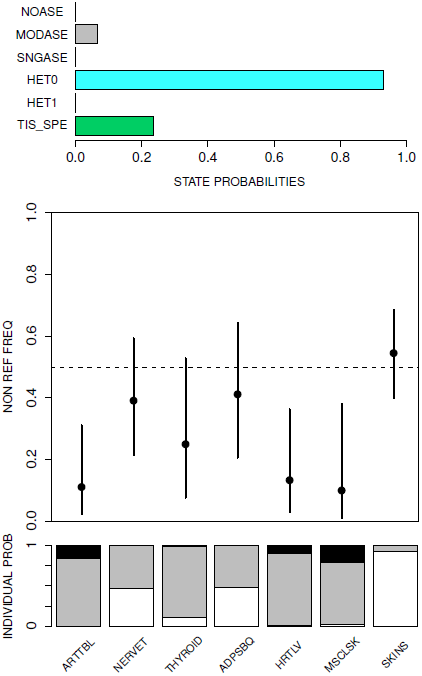
Data on rs121909115. Top panel shows the posterior probabilities for six states as defined in Methods. The tissue specific state (TIS_SPE) is a subset of the heterogeneity states (HET0 and HET1) and the probabilities of the other five states sum to one. Middle panel shows the point estimates of the non-reference allele frequency among RNA-seq reads across seven tissue types (named at the bottom) together with their 95% credible intervals. Bottom panel shows the posterior probability distribution of the group indicator (*γ_s_*) for each tissue type, where white, gray and black denote groups 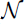, 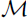 and 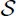, respectively. Tissue types: ARTTBL=Artery tibial, NERVET=Nerve tibial, ADPSBQ=Adipose subcutaneous, HRTLV=Heart ventricle, MSCSKL=Muscle skeletal, SKINS=Skin sun exposed.

The results show that, as expected, most tissue types show ASE where the non-reference allele has lower read counts than the reference allele. In addition, there is evidence of heterogeneity between the tissue (*p*(HET0*|***y**) = 0.93) and that heterogeneity could result from a tissue specific effect where the skin tissue escapes ASE (*p*(TIS_SPE*|***y**) = 0.24). None of the tissues shows strong evidence for complete ASE (group 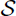), but rather the other six tissue types (apart from skin) are likely to belong to either the moderate ASE group 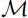, or to no ASE group (nerve and adipose).

The non-reference allele (T) at the SNP rs121909115 is one of the known protein truncating mutations in *ECM1* that in homozygous carriers lead to lipoid proteinosis, also known as Urbach-Wiethe disease, (OMIM 247100), (Hamada *et al.*, 2003). The symptoms of this disease include scarring and infiltration of skin and mucosae (Hamada, 2002). Therefore, it is an interesting observation that in our data the allele with the nonsense mutation is expressed more strongly in the skin tissue than in several other tissue types.

## 4 Discussion

We have introduced a statistical framework to assess similarities and differences in allelic specific expression (ASE) between tissue types. A motivation for our work comes from ongoing RNA-sequencing projects such as the GTEx project (GTEx-Consortium, 2013) that generates data on up to 30 tissue types per individual with read counts per tissue and per site starting from around 10.

We have chosen a Bayesian approach because it leads to a consistent probabilistic quantification of the support that the data provide for each of the competing models. We see this as an advantage over a series of separate analyses, such as, for example, would be needed by an approach that first assessed heterogeneity using the Q-statistic and if no (statistically significant) heterogeneity was observed, would further classify the data set into one of the homogeneous states. Previously, two excellent studies on Bayesian models for expression data have been published by Skelly *et al.* (2011) and Flutre *et al.* (2013).

Skelly *et al.* (2011) consider a three-stage hierarchical model for allele read counts from genome-wide RNA-seq data of one individual. They observe allele counts at each heterozygous SNP (1st level) which are assigned to genes (2nd level) whose common properties are controlled by genome-wide parameters (3rd level). Their model would be directly applicable also to multi-tissue RNA-seq data where tissue specific allele counts replace SNP-specific allele counts and sites replace genes at the second level of the model. Suppose, for example, that we had multi-tissue RNA-seq data on a set of individuals who are heterozygous for at least one protein-truncating variant (PTV). The model of Skelly *et al.* (2011) would produce posterior distribution on the global proportion of the PTVs that show ASE in at least one tissue type, as well as variant specific posterior probability of ASE. However, it would not give as refined characterization of ASE at each PTV as our hierarchical model GTM* (Supplementary Information) and it would not do inference on tissue specific group indicator parameters (*γ_s_*) that allow probabilistic model comparison between different patterns of ASE.

Flutre *et al.* (2013) developed a method for a joint eQTL analysis across multiple tissues. They work with micro-array expression data using a linear model that is not directly applicable to the data sets we have in mind: read count data from several tissues of a single individual. They use tissue specific binary indicator parameters that tell whether a variant is an eQTL in each tissue and introduce three ways to assign prior probabilities to different configurations of the indicators. Their “lite” model gives positive prior on only those configurations whose distance from homogeneity is at most 1. Their “BMA” model is similar to what we have used in that the prior of a configuration depends only on its distance from homogeneity and that the total prior probability corresponding to each value of distance is the same. Finally, their most complex “BMA-HM” model treats the weights of the configurations as random variables and estimates them across genes using a hierarchical model. Similar hierarchical model, that would learn joint ASE patterns between tissues by a simultaneous analysis of a set of genetic variants (e.g. PTVs) is also an important topic for further development of our hierarchical model GTM*.

Even though hierarchical models, such as our GTM* and those of Skelly *et al.* (2011) and Flutre *et al.* (2013), are conceptually attractive, we believe that GTM’s ability to analyze one PTV at a time has its merits from a practical point of view: it is quick to run, easy to understand and requires read count data on only one variant.

### Heterogeneity measures

Our model assumes that all tissues in a same group have the same reference allele read frequency *θ*. In practice, we expect that our model is robust to some heterogeneity within each group, since the priors for different groups are so clearly separable from each other (Fig. 1 and Supp Fig. 1). Our main interest is to assign tissue types into (three) broad categories of ASE, and consequently we call a data set heterogeneous only if some pair of tissues show fairly strong difference in the non-reference allele frequency. Standard heterogeneity measures, such as *Q*, ask a slightly different question of whether parameters *θ_s_* have exactly the same value across the tissues. A frequentist answer to this latter question is given by an empirical P-value of heterogeneity measures *Q* or *p*(HET*|***y**) estimated under the null hypothesis that all tissues have the same value for *θ* parameter which is estimated from the data. By this approach, heterogeneity P-values in the *ECM1* example of Figure 4 are 0.011 and 0.006 by using *Q* and *p*(HET0*|***y**), respectively, as a test statistic. While these P-values point to general heterogeneity between the tissue types, our GTM analysis leads to more detailed information considering the type of heterogeneity: we see heterogeneity where at least one tissue type escapes ASE (our HET0 state).

Our heterogeneity probabilities are based on a lower bound of the true marginal likelihoods of the heterogeneity states. We showed that in a large majority of our data sets (with 10 tissues) the approximation is accurate, but the approximation may not always work as well with larger number of tissues. Therefore, in addition to the heterogeneity probabilities, we also compute, for each pair of tissues, a posterior expectation of the distance between them. Here we define the distance between two tissues to be 0 if they belong to a same group and 1 otherwise. Maximum of all pairwise distances gives an indication whether there is heterogeneity between tissues. By further defining the distance between the ASE groups 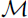 and 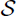 to be 0, we have a measure for particular kind of heterogeneity (HET0) where at least one tissue belongs to the no ASE group 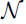.

### 4.1 Application of the Method

**Nonsense-mediated decay.** We envisage that a primary application of our method will be in analyzing nonsense-mediated decay (NMD). Protein truncating variants are usually subject to NMD, a cellular mechanism that detects premature termination codons and prevents expression of truncated proteins. Integrated genome and transcriptome sequencing studies in lymphoblastoid cell lines have demonstrated that allele specific expression can be used for testing variants predicted to trigger NMD (Montgomery *et al.*, 2010; MacArthur *et al.*, 2012; Montgomery *et al.*, 2011; Lappalainen *et al.*, 2013). To test whether the predicted NMD truly happens, we can use the one-sided ASE models rather than the two-sided ones to explicitly require that an ASE signal is present only if the minor allele, and not the major allele, is silenced. We applied the one-sided approach in our example analysis of a PTV in *ECM1*.

**Imprinting.** Genomic imprinting is a phenomenon where only an allele inherited from the parent of a particular sex is expressed (Babak *et al.*, 2008). To assess imprinting for one tissue type one could consider a common coding variant of the gene of interest in that tissue across multiple heterozygous individuals. If the gene is imprinted, then in about half of the individuals the allele 1 should be silenced, because we expect that in about half of the individuals the allele 1 at this locus is inherited from the mother and in the other half it originates from the father. We could extend our framework to such a case by allowing each ASE group to be divided into two subgroups with proportion parameters *θ* and 1*−θ*, respectively, where *θ* has a Beta-prior as in our GTM model.

**Modest ASE effects.** An interesting application of allele specific expression is in the study of *cis*-regulatory variants in LD with coding variants. When regulatory variants have only modest effects (Dimas *et al.*, 2009), we could modify our prior on ASE states to reflect this. With this approach we envision that researchers are able to study the effect of *cis*-regulatory variants on transcription across a broad range of tissues where the number of samples per tissue may be limited. However, compared to strong ASE effects, more modest effects require much larger read counts per tissue and decrease the ratio between biological signal and possible technical noise (Degner *et al.*, 2009).

**Multiple individuals.** Suppose we have RNA-seq data on the same tissue types from several individuals who are heterozygous at a particular variant. We could first assess, for each tissue type, whether the individuals are heterogeneous in their ASE status. If there is no evident heterogeneity, a simple approach is to combine the reads from the same tissue type across the individuals before analyzing the data across the tissues. A more refined model that accounts for possible individual specific effects that are shared across the tissues requires further work.

## 5 Conclusion

We have presented grouped tissue model (GTM) and its multi-site extension (GTM*), to (i) classify tissues into three groups at each site according to their allele specific expression (ASE) patterns and (ii) classify the sites into five combined ASE states according to their tissue-wide ASE profiles. We see major applications of our approach in assessing homogeneity, heterogeneity and tissue specificity across a group of genetic variants assumed to have similar properties, e.g. variants predicted to trigger nonsense-mediated decay, variants in genes with evidence of imprinting, or variants in LD with GWAS loci for a particular disease.

As an example, we presented an application of the method to read count data from the GTEx project for one heterozygote carrier of a premature truncating mutation (p.R53X, rs121909115) in the *ECM1* gene. For this variant, which in homozygous form is known to cause lipoid proteinosis, we identified heterogeneous gene expression effects across tissues and evidence for a complete escape from nonsense-mediated decay in skin tissue.

The identification and characterization of ASE, in particular for protein truncating variants with a putative complete loss of function effect, will provide a better understanding of the biological mechanisms that are involved in transcriptional regulation and will improve our computational models for annotating variants identified in case-control or clinical genome sequencing studies.

## Acknowledgement

We thank members of the GTEx Consortium for useful comments; the sequencing platforms at the Broad Institute for generating the DNA and RNA sequencing data; the Biospecimen Source Sites (BSS) for collecting the samples and the Laboratory, Data Analysis, and Coordinating Center (LDACC) for handling, performing quality control, and processing the data.

## References

1. Babak, T., DeVeale, B., Armour, C., Raymond, C., Cleary, M. A., van der Kooy, D., Johnson, J. M., and Lim, L. P. (2008). Global survey of genomic imprinting by transcriptome sequencing. Current biology, 18(22), 1735–1741.

2. Degner, J. F., Marioni, J. C., Pai, A. A., Pickrell, J. K., Nkadori, E., Gilad, Y., and Pritchard, J. K. (2009). Effect of read-mapping biases on detecting allele-specific expression from RNA-sequencing data. Bioinformatics, 25(24), 3207–3212.

3. Dimas, A. S., Deutsch, S., Stranger, B. E., Montgomery, S. B., Borel, C., Attar-Cohen, H., Ingle, C., Beazley, C., Arcelus, M. G., Sekowska, M., et al. (2009). Common regulatory variation impacts gene expression in a cell type–dependent manner. Science, 325(5945), 1246–1250.

4. Flutre, T., Wen, X., Pritchard, J., and Stephens, M. (2013). A statistical framework for joint eQTL analysis in multiple tissues. PLoS genetics, 9(5), e1003486.

5. Grundberg, E., Small, K. S., Hedman, Å. K., Nica, A. C., Buil, A., Keildson, S., Bell, J. T., Yang, T.-P., Meduri, E., Barrett, A., et al. (2012). Mapping cis-and trans-regulatory effects across multiple tissues in twins. Nature genetics, 44(10), 1084–1089.

6. GTEx-Consortium (2013). The genotype-tissue expression (GTEx) project. Nature genetics, 45(6), 580–585.

7. Gutierrez-Arcelus, M., Lappalainen, T., Montgomery, S. B., Buil, A., Ongen, H., Yurovsky, A., Bryois, J., Giger, T., Romano, L., Planchon, A., Falconnet, E., Bielser, D., Gagnebin, M., Padioleau, I., Borel, C., Letourneau, A., Makrythanasis, P., Guipponi, M., Gehrig, C., Antonarakis, S. E., and Dermitzakis, E. T. (2013). Passive and active DNA methylation and the interplay with genetic variation in gene regulation. eLife, 2.

8. Hamada, T. (2002). Lipoid proteinosis. Clinical and experimental dermatology, 27(8), 624–629.

9. Hamada, T., Wessagowit, V., South, A. P., Ashton, G. H. S., Chan, I., Oyama, N., Siriwattana, A., Jewhasuchin, P., Charuwichitratana, S., Thappa, D. M., Lenane, P., Krafchik, B., Kulthanan, K., Shimizu, H., Kaya, T. I., Erdal, M. E., Paradisi, M., Paller, A. S., Seishima, M., Hashimoto, T., and McGrath, J. A. (2003). Extracellular matrix protein 1 gene (ECM1) mutations in lipoid proteinosis and genotype-phenotype correlation. Journal of investigative dermatology, 120, 34–350.

10. Lappalainen, T., Sammeth, M., Friedlander, M. R., /‘t Hoen, P. A. C., Monlong, J., Rivas, M. A., Gonzalez-Porta, M., Kurbatova, N., Griebel, T., Ferreira, P. G., Barann, M., Wieland, T., Greger, L., van Iterson, M., Almlof, J., Ribeca, P., Pulyakhina, I., Esser, D., Giger, T., Tikhonov, A., Sultan, M., Bertier, G., MacArthur, D. G., Lek, M., Lizano, E., Buermans, H. P. J., Padioleau, I., Schwarzmayr, T., Karlberg, O., Ongen, H., Kilpinen, H., Beltran, S., Gut, M., Kahlem, K., Amstislavskiy, V., Stegle, O., Pirinen, M., Montgomery, S. B., Donnelly, P., McCarthy, M. I., Flicek, P., Strom, T. M., Consortium, T. G., Lehrach, H., Schreiber, S., Sudbrak, R., Carracedo, A., Antonarakis, S. E., Hasler, R., Syvanen, A.-C., van Ommen, G.-J., Brazma, A., Meitinger, T., Rosenstiel, P., Guigo, R., Gut, I. G., Estivill, X., and Dermitzakis, E. T. (2013). Transcriptome and genome sequencing uncovers functional variation in humans. Nature, 501(7468), 506–511.

11. MacArthur, D. G., Balasubramanian, S., Frankish, A., Huang, N., Morris, J., Walter, K., Jostins, L., Habegger, L., Pickrell, J. K., Montgomery, S. B., Albers, C. A., Zhang, Z. D., Conrad, D. F., Lunter, G., Zheng, H., Ayub, Q., DePristo, M. A., Banks, E., Hu, M., Handsaker, R. E., Rosenfeld, J. A., Fromer, M., Jin, M., Mu, X. J., Khurana, E., Ye, K., Kay, M., Saunders, G. I., Suner, M. M., Hunt, T., Barnes, I. H., Amid, C., Carvalho-Silva, D. R., Bignell, A. H., Snow, C., Yngvadottir, B., Bumpstead, S., Cooper, D. N., Xue, Y., Romero, I. G., Wang, J., Li, Y., Gibbs, R. A., McCarroll, S. A., Dermitzakis, E. T., Pritchard, J. K., Barrett, J. C., Harrow, J., Hurles, M. E., Gerstein, M. B., and Tyler-Smith, C. (2012). A systematic survey of loss-of-function variants in human protein-coding genes. Science, 335(6070), 823–8.

12. McCarroll, S. A., Huett, A., Kuballa, P., Chilewski, S. D., Landry, A., Goyette, P., Zody, M. C., Hall, J. L., Brant, S. R., Cho, J. H., et al. (2008). Deletion polymorphism upstream of IRGM associated with altered IRGM expression and Crohn’s disease. Nature genetics, 40(9), 1107–1112.

13. Montgomery, S. B., Sammeth, M., Gutierrez-Arcelus, M., Lach, R. P., Ingle, C., Nisbett, J., Guigo, R., and Dermitzakis, E. T. (2010). Transcriptome genetics using second generation sequencing in a Caucasian population. Nature, 464(7289), 773–777.

14. Montgomery, S. B., Lappalainen, T., Gutierrez-Arcelus, M., and Dermitzakis, E. T. (2011). Rare and common regulatory variation in population-scale sequenced human genomes. PLoS genetics, 7(7), e1002144.

15. Rivas, M. A., Beaudoin, M., Gardet, A., Stevens, C., Sharma, Y., Zhang, C. K., Boucher, G., Ripke, S., Ellinghaus, D., Burtt, N., et al. (2011). Deep resequencing of GWAS loci identifies independent rare variants associated with inflammatory bowel disease. Nature genetics, 43(11), 1066–1073.

16. Rivas, M. A., Pirinen, M., Neville, M. J., Gaulton, K. J., Moutsianas, L., Lindgren, C. M., Karpe, F., McCarthy, M. I., and Donnelly, P. (2013). Assessing association between protein truncating variants and quantitative traits. Bioinformatics, 29(19), 2419–2426.

17. Ronald, J., Akey, J. M., Whittle, J., Smith, E. N., Yvert, G., and Kruglyak, L. (2005). Simultaneous genotyping, gene-expression measurement, and detection of allele-specific expression with oligonucleotide arrays. Genome research, 15(2), 284–291.

18. Skelly, D. A., Johansson, M., Madeoy, J., Wakefield, J., and Akey, J. M. (2011). A powerful and flexible statistical framework for testing hypotheses of allele-specific gene expression from rna-seq data. Genome Research, 21(10), 1728–1737.

19. Sun, W. (2012). A statistical framework for eQTL mapping using RNA-seq data. Biometrics, 68(1), 1–11.

20. Zhang, K., Li, J. B., Gao, Y., Egli, D., Xie, B., Deng, J., Li, Z., Lee, J.-H., Aach, J., Leproust, E. M., et al. (2009). Digital RNA allelotyping reveals tissue-specific and allele-specific gene expression in human. Nature methods, 6(8), 613–618.

21. Zuk, O., Schaffner, S. F., Samocha, K., Do, R., Hechter, E., Kathiresan, S., Daly, M. J., Neale, B. M., Sunyaev, S. R., and Lander, E. S. (2014). Searching for missing heritability: Designing rare variant association studies. Proceedings of the National Academy of Sciences, 111(4), E455–E456.

